# A Cre-dependent massively parallel reporter assay allows for cell-type specific assessment of the functional effects of non-coding elements *in vivo*

**DOI:** 10.1101/2021.05.17.444514

**Authors:** Tomas Lagunas, Stephen P. Plassmeyer, Anthony D. Fischer, Ryan Z. Friedman, Michael A. Rieger, Din Selmanovic, Simona Sarafinovska, Alessandra F. Aguilar Lucero, Joon-Yong An, Stephan J. Sanders, Barak A. Cohen, Joseph D. Dougherty

**Affiliations:** Department of Genetics, Washington University School of Medicine, 660 S. Euclid Ave, Saint Louis MO, 63108, USA; Department of Psychiatry, Washington University School of Medicine; Division of Biology and Biomedical Sciences, Washington University School of Medicine; Department of Psychiatry and Behavioral Sciences, UCSF Weill Institute for Neuroscience, University of California, San Francisco, CA 94518; Department of Integrated Biomedical and Life Science, Korea University, Seoul 02841, Republic of Korea; School of Biosystem and Biomedical Science, College of Health Science, Korea University, Seoul 02841, Republic of Korea

**Author notes:** **Contact Information:** Dr. Joseph Dougherty, Department of Genetics, Campus Box 8232, 4566 Scott Ave., St. Louis, MO. 63110-1093, E.

## Abstract

The function of regulatory elements is highly dependent on the cellular context, and thus for understanding the function of elements associated with psychiatric diseases these would ideally be studied in neurons in a living brain. Massively Parallel Reporter Assays (MPRAs) are molecular genetic tools that enable functional screening of hundreds of predefined sequences in a single experiment. These assays have not yet been adapted to query specific cell types *in vivo* in a complex tissue like the mouse brain. Here, using a test-case 3′UTR MPRA library with genomic elements containing variants from ASD patients, we developed a method to achieve reproducible measurements of element effects *in vivo* in a cell type-specific manner, using excitatory cortical neurons and striatal medium spiny neurons as test cases. This new technique should enable robust, functional annotation of genetic elements in the cellular contexts most relevant to psychiatric disease.

## Introduction

In the current era of common and rare variant genome-wide approaches, thousands of candidate genetic variants with potential association to psychiatric and neurological diseases have been uncovered, the vast majority in noncoding, presumably regulatory, DNA elements. For common variants, large collaborative studies have identified dozens of genomic regions that are significantly associated with disease ^1–3^, but each region contains hundreds to thousands of elements containing noncoding variants, of which only a subset are thought to have a functional consequence and potentially be causal. For rare variants, whole-genome sequencing has identified thousands of noncoding variants per individual, and efforts at associating these with disease would benefit from knowing which are found in elements that control gene expression in the brain, and thus might alter neuronal function. However, in either case, defining the effect of DNA elements has proven to be a major challenge given the large number that need to be screened. Furthermore, cell-type context plays an important role in gene-regulation studies ^4^. For example, as they mature, neurons express a variety of neuron-specific transcription factors (TFs) (e.g. ^5^) and RNA-binding proteins (RBPs) ^6^, and thus elements containing their binding sites for these would only show effects in mature neurons. Therefore, there is a need for a high-throughput method that can be easily adapted to functionally screen elements in a parallel fashion, specifically in the cellular contexts relevant to diseases of the central nervous system (CNS). For most psychiatric diseases, this ideal cellular context would be specific classes of neurons, *in vivo*.

In the past decade, numerous *de novo* mutations have been directly implicated in ASD ^7,8^. Initial analyses focused on mutations in coding regions, which are more readily interpreted for functional effects than noncoding variants ^9–13^. However, there is estimated to be substantial additional burden from noncoding mutations ^14,15^. This can include both transcriptional regulators, like promoters and enhancers, as well as 5′/3′ untranslated regions (UTRs). UTRs contain several classes of regulatory elements that control mRNA stability, subcellular localization, and rate of translation for their cognate transcript ^16^. However, these regions pose challenges to study for functional effects since they don’t follow a triplet code and are not easily interpretable.

Massively Parallel Reporter Assays (MPRAs) are genetic tools that could address these challenges since they can be used to functionally assay several thousand predefined sequences at once ^4^. These assays have enabled functional annotation of thousands of noncoding genomic elements, as well as the impact of variants in UTRs in particular, prioritizing potentially causal changes ^17–20^. In addition, recent MPRA studies have begun to dissect the role of 3′UTR variation ^21^ in function and regulatory activity *in vitro*. Unsurprisingly, there is only a modest overlap of functional elements across six diverse human cell lines, underscoring the density of elements with cell type-specific regulatory potential within UTRs. Furthermore, there are limits to the extent to which an *in vitro* system, even primary cells or iPSC derived neural systems, can recapitulate the normal gene expression and thus regulatory landscape seen during neuronal development *in vivo*. Thus, in the context of neuropsychiatric disease, elements would ideally be assayed in the brain and in relevant cell types in order to more accurately model the effect of these variants.

Here, we describe the development of a high-throughput cell-type specific MPRA approach for the mouse brain, with the sensitivity to measure the effects of individual elements, using a Cre recombinase-dependent library design. As a test-case, we used a 3′UTR MPRA library to functionally assay several hundred elements containing *de novo* variants found in the genomes of ASD cases and sibling controls. We first piloted this in a mouse neuroblastoma cell line, assessing total RNA and RNA paired with a ribosome affinity purification to enable assessment of both transcriptional and translational effects. We then optimized the delivery of these same elements to two types of neurons *in vivo*. We were indeed able to assess the functional effects of hundreds of elements in parallel, and found effects of elements are highly cell type-specific. We also examined the ability to test for variant effects, and power calculations indicate this should be possible, but will require more extensive barcoding than used here. In all, the approach here should enable future large-scale assessment of the functional impact of variants from psychiatric genetics in specific cell types in the brain.

## Results

### Cre-dependent MPRA reproducibly measures element effects in a mouse neuroblastoma cell line

As a proof of principle, we examined *de novo* variants identified within annotated 3′UTRs from the whole-genome sequencing of 519 families with ASD, primarily from the Simons Simplex Collection ^8^, targeting 342 mutations from probands and 307 from unaffected siblings within the same cohort (649 unique variants [**Supp. Table 1]** to make 1298 ref/alt pairs). For each variant we synthesized an allelic pair of 3′UTR ‘elements’ spanning 120 bp of sequence centered on the variant. To be able to compare biological to non-biological sequence elements, for 322 variants, we randomly shuffled the sequence to generate a set of GC-matched controls. Additionally, we included 4 predicted stabilizing/destabilizing controls. We tagged all 1,624 elements with six unique barcodes to provide internal replicates and be able to measure potential for barcode effects. To enable eventual cell-type specific studies, we cloned the final library of 9,744 synthesized oligos into the 3′UTR of a membrane-localized tdTomato reporter embedded in a Double-floxed inverse Orientation (DiO) cassette ^22^, such that the reporter library would only express following Cre-mediated recombination **[Fig. 1A-B]**.

**Fig 1.**
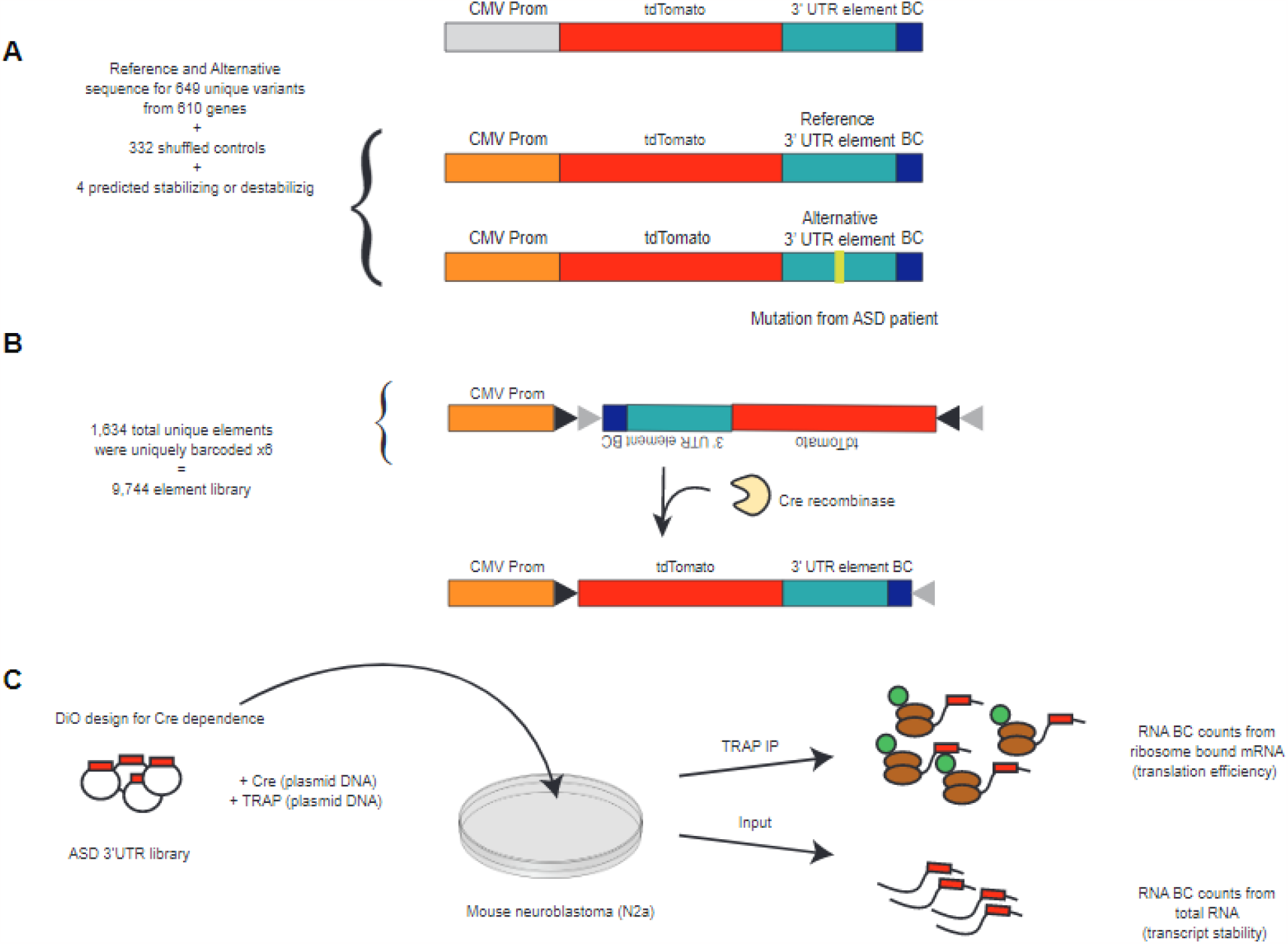
ASD 3′UTR library design and delivery. A) MPRA library constructs were designed with a CMV promoter (prom) driving the TdTomato reporter, followed by the 3′UTR oligo reference or alternative sequence (with or without variant, respectively) that is uniquely barcoded (BC). B) All elements were uniquely barcoded 6 times. Cloning of this library was completed in the Double-floxed inverse Orientation (DiO) design for Cre-dependent expression. C) MPRA library, Cre recombinase and TRAP components were delivered via plasmid transfection into N2a cells. Following incubation, total RNA and TRAP RNA were isolated to prepare sequencing libraries to then count BCs to calculate expression per element.

To first evaluate whether our assay could detect UTR element effects on reporter transcript abundance and translation, we co-transfected the library into mouse neuroblastoma N2a cells with two additional constructs–one expressing Cre recombinase, and another expressing eGFP-tagged large ribosomal subunit protein L10a (eGFP-RPL10a). The eGFP-RPL10a construct allows us to employ the Translating Ribosome Affinity Purification (TRAP) technique to measure the effects of UTR elements on ribosome occupancy ^23^. We harvested RNA from six replicate transfections from both the whole-cell lysate (Input) and the polysome-bound TRAP fraction ^24^ **[Fig. 1C]**. Barcode sequencing libraries were prepared from both Input RNA and TRAP RNA to identify elements that alter ribosome occupancy (TRAP) on top of effects on transcript abundance (Input). We also conducted DNAseq on the plasmid DNA re-extracted from the transfected cells to enable normalization of each RNA barcode to its starting abundance in the cells.

We examined the coverage and reproducibility of the assay, and the range of the biological activity across elements. We sequenced to an average depth of 5,388 counts per barcode. In the DNA, 8,053 barcodes had non-zero counts, suggesting a <20% element dropout at the cloning stage. Cloning efficiency correlated with element GC-content, as elements with less than 40% GC content cloned less efficiently **[Supp. Fig. 1A]**. A corresponding 85% of elements were represented with at least three barcodes and carried forward for analysis **[Supp. Fig. 1B]**. In the RNA data, correlations of barcode abundance between replicate libraries from both Input and TRAP generally exceeded 0.99 (Pearson’s Correlation Coefficient, PCC) **[Fig. 2A]**, indicating high reproducibility (read depth, correlations, etc, for all experiments in paper are summarized in **Supp. Table 2**). Correlations of either RNA measure with barcode abundance in recovered plasmid libraries averaged 0.99 (PCC), indicating that variation in reporter abundance was largely driven by DNA copy number, as the range of differences in cloning efficiency exceeds the magnitude of expected biological effects of elements. Thus, we normalized input RNA counts to plasmid DNA counts for subsequent analyses. This revealed variation in steady state RNA abundance across elements, with 99% of elements spanning -1.76 to 1.14 log_2_-normalized expression (RNA/DNA) **[Fig. 2B]**, indicating that the sampled UTR elements exhibit a 7-fold range in transcript abundance as measured by our assay.

**Fig 2.**
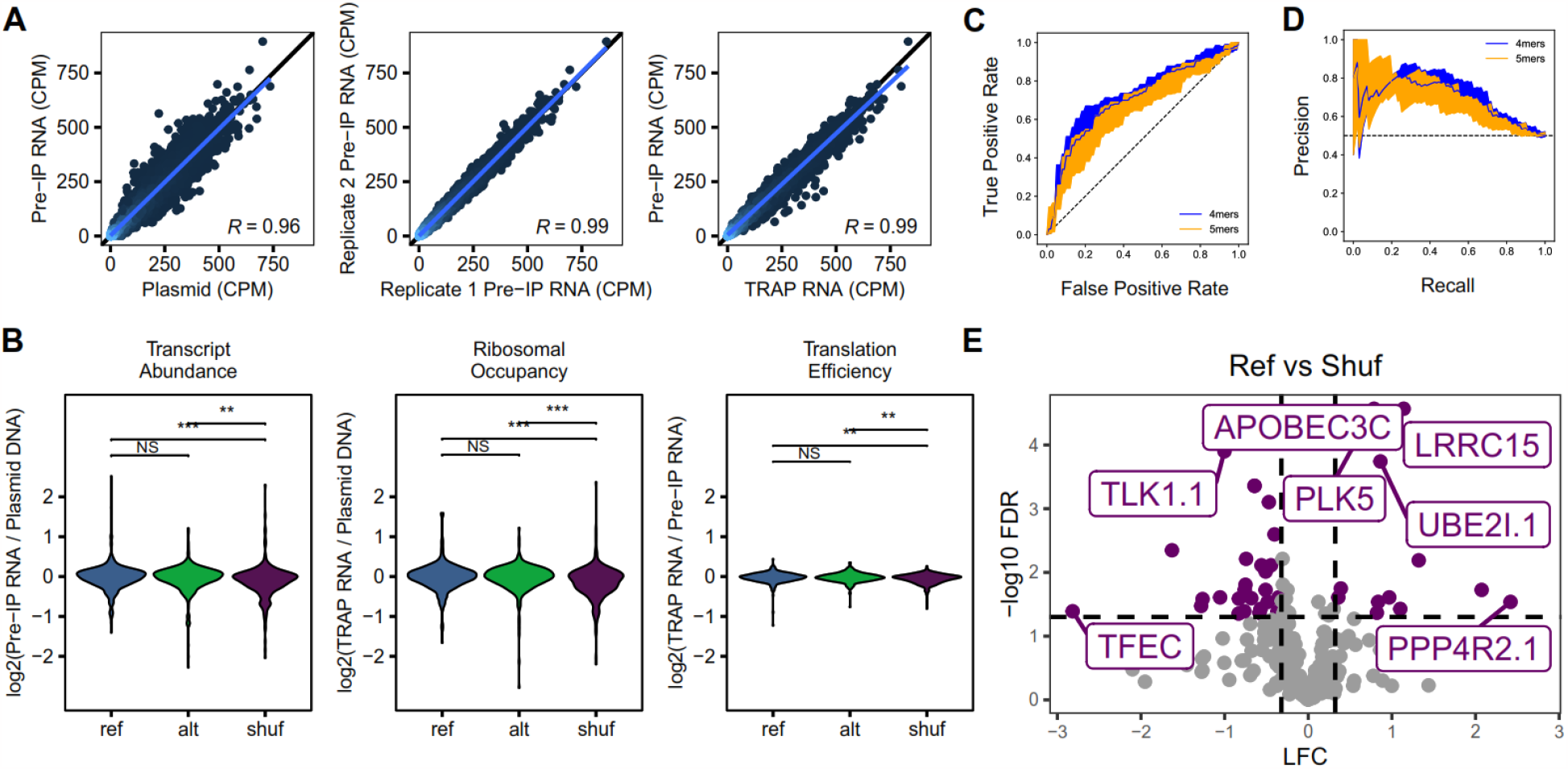
Screen in mouse neuroblastoma cell line identifies variants that alter steady state transcript abundance. A) Scatter plots showing correlation between replicates of RNA vs RNA (left), RNA vs plasmid DNA (center), and RNA vs TRAP RNA (right) CPM counts. B) Pairwise comparison of expression value distribution among Ref, Alt, and Shuf sequences in Transcript Abundance, Ribosomal Occupancy, and Translation Efficiency data sets. Significance is denoted by asterisk. C) ROC and D) PRC curves for k-mer SVMs to classify high and low expressing elements. Shaded area represents 1 standard deviation based on five-fold cross-validation E) Volcano plot for Ref vs Shuf elements (purple) in library showing significance (y-axis) vs log2 FC (x-axis). Horizontal dashed line corresponds to FDR 0.05 and vertical dashed lines correspond to log2 FC equivalent to 25% change in expression. Full results list can be found in Supplement Table 3, worksheet 1.

Normalizing TRAP RNA abundance by DNA copy number revealed a similar dynamic range in the ribosomal occupancy of reporter transcripts. However, these differences are driven primarily by the underlying difference in transcript abundance. Normalizing TRAP RNA abundance to the input RNA abundance a proxy for ‘Translation Efficiency’ (TE), defined here as log_2_ TRAP/Input counts, showed a narrow dynamic range from -0.46 to 0.29, indicating 3′UTR element effects on translation regulation are more subtle than on transcript abundance. Interestingly, one novel observation arose from a pairwise comparison of genome-derived reference elements to GC-matched shuffled control sequences. Specifically, random sequences had both lower transcript abundance (Wilcoxon signed-rank p = 1.69×10^−4^) and TE (p = 1.67×10^−4^) than their corresponding reference sequences **[Fig. 2B]**. This suggests that genomic sequences generally promote higher steady-state transcript abundance and ribosome occupancy than random sequences. However, the elements containing *de novo* variants (alternative alleles; Alt) did not show a systematic difference from their paired reference allele (Ref) elements. This is not unexpected, as most are small or single base mutations, and only a small subset of human mutations, even from probands, might be presumed to be strongly functional *a priori*, and the effect of the alternate allele might be in either direction.

Biological effects should be driven by specific sequence elements in the UTRs, and thus activity should be somewhat predictable from primary sequence. To establish a biological signature of elements that increase or decrease RNA levels, we trained k-mer support vector machines (SVMs) ^25^ to classify the 200 highest-expressing elements from the 200 lowest-expressing elements, pooling Ref and shuffled (Shuf) sequences. In this framework, each sequence is represented by the frequency of all possible k-mers as input to the SVM. We trained 4- and 5-mer SVMs with 5-fold cross-validation. To ensure the SVM was not overfit, we also fit SVMs on the same sequences with random labels. The SVMs achieved an area under the receiver operating characteristic (AUROC) of between 0.707 and 0.663, for 4-mer and 5-mer models respectively, and an area under the precision recall curve (AUPRC) between 0.693 and 0.669 for these same models **[Fig. 2C-D]**. Models fit on random labels could not classify the data (AUROC between 0.486 and 0.521) **[Supp. Fig. 2]**, indicating there are sequence-specific elements underlying UTR activity. To understand which sequences mediated these effects, we next scored all possible 4-mers against the 4-mer SVM. 4-mers predicted to be highly active tended to be GC rich, while 4-mers predicted to be inactive tended to be AT rich. We also used DREME ^26^ to identify *de novo* motifs enriched in the high expressing sequences relative to the low expressing sequences and obtained similar results. Taken together, these results indicate a substantial fraction of the activity of UTRs is driven by sequence features captured by small motifs, and identifies the motifs with activity in N2a cells. It also indicates that for the effect sizes studied here, any effects of individual barcodes on transcript abundance were sufficiently small to not obscure the consistent biology detected by the SVM.

While more highly expressed elements tended to be GC rich, genomic elements were clearly different from random GC matched controls. Comparing each Ref element to its matched Shuf control revealed that 54 were significantly different (Benjamini-Hochberg FDR <0.05) with a median 1.65-fold change in expression **[Fig. 2E, Supp. Table 3]**. Thus, genomic sequences produce a specific level of activity upon which allelic effects are expected to act. Of the 303 tested comparisons, 40 showed both a significant difference and a >25% magnitude change in expression. Of the significant changes, 38 were downregulating. Assuming equal probability of up- and down-regulation, this is more than expected by chance (hypergeometric p = 0.0033, OR = 1.69), again reflecting the relative greater propensity for genomic derived UTR tiles to enhance steady-state reporter expression. Contrasting effects on transcript abundance with ribosome occupancy again revealed that variant effects on TE tended to be much smaller. Overall, our cell line assay confirmed reproducibility and robustness of our measurements of element level activity by our DiO 3′UTR MPRA design, motivating applying the approach to specific cell types *in vivo*.

### Cre-dependent MPRA reproducibly measures functional effects of several hundred UTR elements in excitatory neurons in the mouse brain

To assess the effect of these elements *in vivo*, the entire element library was packaged in adeno-associated virus serotype 9 (AAV9) for delivery into the mouse brain. We have previously shown ^27^ that with AAV9 delivery we get widespread viral transduction in the neocortex and mainly target neurons and astrocytes. We found that packaging of the library did not drastically change the range of distribution or barcode recovery rates and correlated well (PCC > 0.8) with the plasmid counts **[Supp. Fig 3]**. Thus, as packaging had no adverse effects on the composition of the library we moved forward with delivery *in vivo*.

Bioinformatic analysis of the expression patterns of genes associated with ASD have revealed a correlation structure of two loose modules - a module enriched for chromatin regulators with peak expression in immature excitatory neurons, and a module of synaptic-related proteins, with peak expression during critical periods of postnatal synaptogenesis and pruning ^7,28–30^. Therefore, we first attempted to deliver the library to two neuronal subtypes, layer V pyramidal neurons and cortical GABAergic interneurons, during this pruning period by using *Rbp4* and *Vgat* Cre driver lines, respectively. However, especially when using an untargeted P1 injection strategy, we discovered that only a small fraction of the delivered elements were recovered, the representation of barcodes was highly distorted and, in many cases, favoring a small, distinct subset in each biological replicate, resulting in low correlation between replicates (PCC < 0.2). This could either be due to high biological variability or a technical effect termed ‘jackpotting’ which here we define as having barcodes whose final measurement in the MPRA sequencing library does not reflect their starting abundance in the sample RNA likely because only a subset of the barcodes were sampled at a particular step in the library preparation. We conducted extensive experiments to determine at which step such jackpotting might occur (see methods), and traced it to the very first step. This indicated that having the Cre recombinase only in a small fraction of the cells resulted in a very low ‘library density in the total RNA, meaning only a very small fraction of the RNA molecules in the sample came from the MPRA library.

Since *Rbp4-positive* and *Vgat*-positive cells made up a small population of cells in the mouse brain, to increase the library density we delivered the AAV library to a well-characterized excitatory neuron specific Cre line (Vglut1-IRES2-Cre-D^31^; Vlgut1^Cre^) at P0-P2 **[Fig. 3A]**, which makes up a larger population of cells, covering all pyramidal cells of the cortex. We first confirmed the expression of the library by immunofluorescence **[Fig. 3B]**. We saw widespread expression of the tdTomato reporter in cells with the morphology of pyramidal neurons with the perinatal injection yielding transductions across cortex. Importantly, Cre negative littermates showed no expression of the library, confirming cell-type specificity **[Fig. 3D]**. Next, an additional 11 animals’ (6 males and 5 females) cortices were collected for RNA at P21. We sequenced, in all, 11 RNA replicates and 2 replicates of viral prep DNA to obtain RNA barcode and DNA barcode counts, respectively.

**Fig 3.**
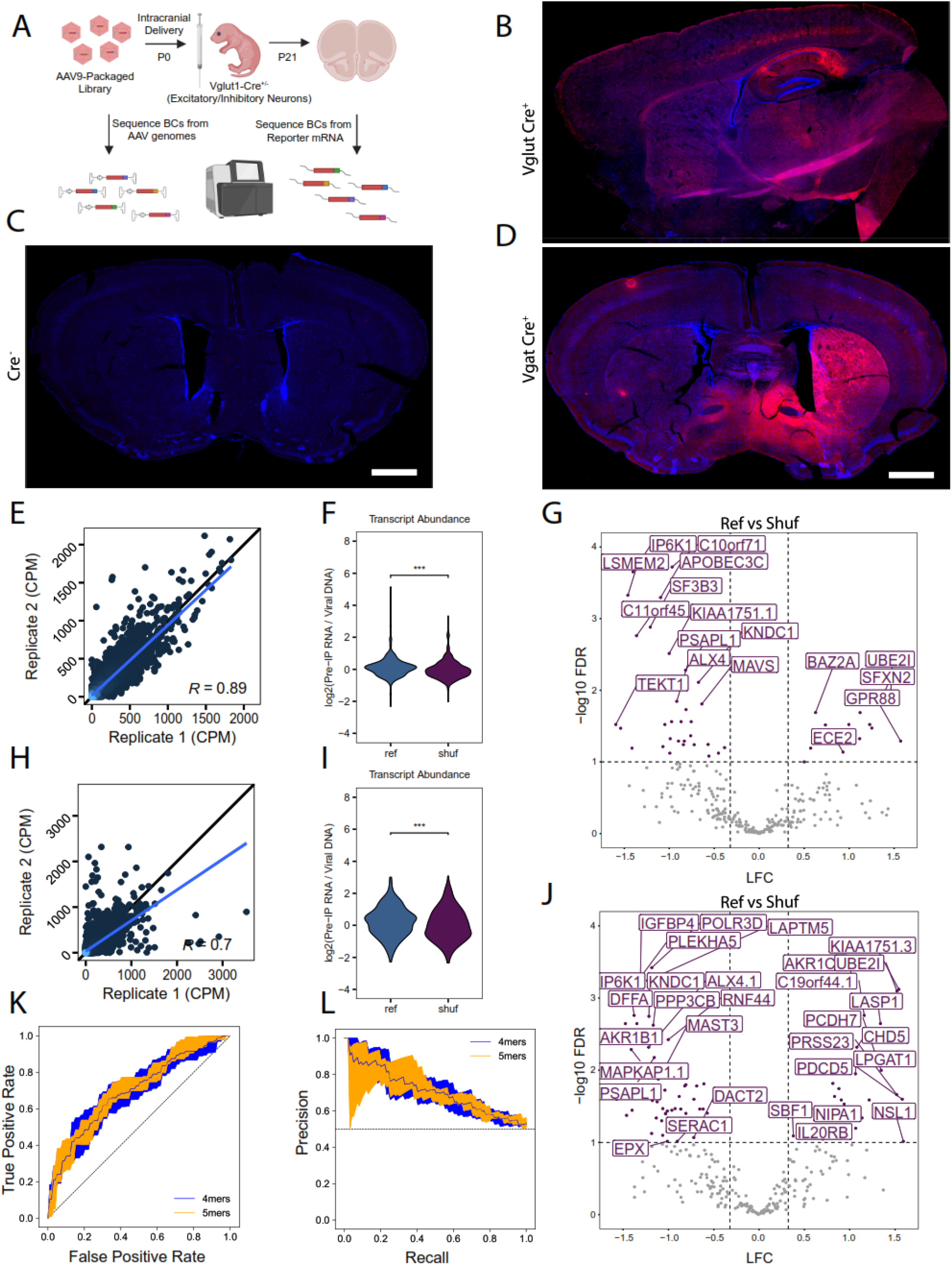
Screen in excitatory neurons in the mouse brain identifies elements that alter steady state transcript abundance. A) MPRA library was packaged into AAV9 and delivered into perinatal mouse cortices via intracranial injection and later harvested at P21 for RNA extraction. Libraries were prepared from AAV genomes and reporter mRNA BCs. B) Immunofluorescence of P21 brain showing localization of tdTomato expression (from MPRA library) in excitatory neurons in cortex with nuclei labeled with DAPI (blue). C) Immunofluorescence of Cre-negative littermate showing DAPI staining and no signal from the Cre-dependent MPRA library. D) Immunofluorescence of P21 brain showing localization of tdTomato expression (from MPRA library) in VGAT+ spiny neurons of striatum. E) Scatter plot showing correlation between RNA counts of biological replicates in excitatory neurons. F) pairwise comparison of Ref, and Shuf sequence expression in excitatory neurons. G) Volcano plot for Ref vs Shuf element expression in excitatory neurons in library showing significance (y-axis) vs log2 FC (x-axis). Horizontal dashed line corresponds to FDR 0.05 and vertical dashed lines correspond to log2 FC equivalent to 25% change in expression. Full results list can be found in Supplement Tables 3 and 4, worksheet 3. H) Scatter plot showing correlation between RNA counts of biological replicates in medium spiney neurons. I) pairwise comparison of Ref, and Shuf sequence expression in medium spiney neurons. J) Volcano plot for Ref vs Shuf element expression in medium spiney neurons. Full results list can be found in Supplement Tables 3 and 4, worksheet 4. K) ROC and L)PRC curves for k-mer SVMs to classify high and low expressing elements in excitatory neurons. Shaded area represents 1 standard deviation based on five-fold cross-validation.

Next, we performed a similar quality control analysis as for N2a data above. Correlations of barcode abundance between biological replicates on average exceeded 0.80 (PCC) **[Fig. 3E]**. Notably, this observed correlation is lower than our *in vitro* test, but increased variability is commensurate with lower rates of element delivery and recovery from a subset of cells in complex tissue. (This increased variance also motivated our doubling of the number of replicates *in vivo* relative to *in vitro*). Similar to what was done for the N2a data, we removed elements which were absent in the DNA counts and filtered for a minimum sequencing depth and barcode number, resulting in 313 analyzed elements. Pairwise comparison of genome-derived Ref elements to GC-matched Shuf control sequences again showed that Shuf sequences had lower transcript abundance (Wilcoxon signed-rank p = 2.99×10^−4^) than their corresponding Ref sequences, as observed in N2as **[Fig. 3F]**. Of the 301 testable Ref-Shuf comparisons, 36 showed a significant difference in expression. While we observed a 25:11 ratio of downregulation:upregulation effect of the reference sequences vs. shuffled sequences, the enrichment was not significant (hypergeometric test, p-value = 0.14, OR = 1.19). **[Fig. 3G, Supp. Table 3]** Finally, we again used k-mer SVMs to determine if there were sequence features that predicted *in vivo* activity and achieved an AUROC between 0.696 and 0.706, for 4-mer and 5-mer models respectively, and an AUPRC between 0.708 and 0.704 for these same models **[Fig. 3K-L]**, comparable to the SVMs trained on *in vitro* activity. Thus, for the large effect sizes observed when changing all sequences in an element, our approach was sufficiently powered.

### In vivo Cre-dependent MPRA works reproducibly in more than one cell type

We next sought to determine if the method was effective across cell types. We returned to the Vgat (GABAergic) mouse line but with a modifications of our prior approach - we focused our delivery and dissection on the striatum, an area that is both of high interest for ASD^32^, but also where >90% of the neurons are GABAergic, making it easier to deliver the library into a large fraction of the of cells. This similarly resulted in better correlations than prior experiments **[Fig. 3H]**, and again genomic sequences generally had greater abundance than shuffled controls [**Fig. 3I]**, with dozens having significant activity **[Fig 3J, Supp. Table 3]**. In all, this demonstrates that the Cre-dependent MPRA is effective across more than one cell type, and that we were well-powered to detect the effect sizes seen when comparing reference sequences to random sequence controls.

Thus, the Cre-dependent MPRA should allow quantification of the impact on transcript abundance of a given element in specific cell types in the brain. This is essential because neurons have vastly different expression of *trans*-acting factors (e.g., TFs, RBPs, miRNAs) than cell lines. This is highlighted by the low correlation of expression values of element activity across N2As compared to excitatory neurons **[Fig. 4A]**. Furthermore, transcript abundance spanned a broader range in the *in vivo* assay (Brown-Forsythe Levene-type p < 2.2×10^−16^), highlighting the possibility that a more complex regulatory environment may contribute to a greater dynamic range **[Fig. 4B]**. Finally, the cross-validated SVM scores of the N2a activity are uncorrelated to the observed activity in excitatory neurons **[Fig. 4C]**, consistent with cell type-specific factors regulating UTR activity through interaction with specific sequences. This highlights the need to assess the function of noncoding elements in multiple contexts, and especially focusing on contexts where noncoding variants for a specific disease are most likely to act. Comparing our two neuronal types together reveals a somewhat better correlation, but with some differences [**Fig 4D]**.

**Figure 4.**
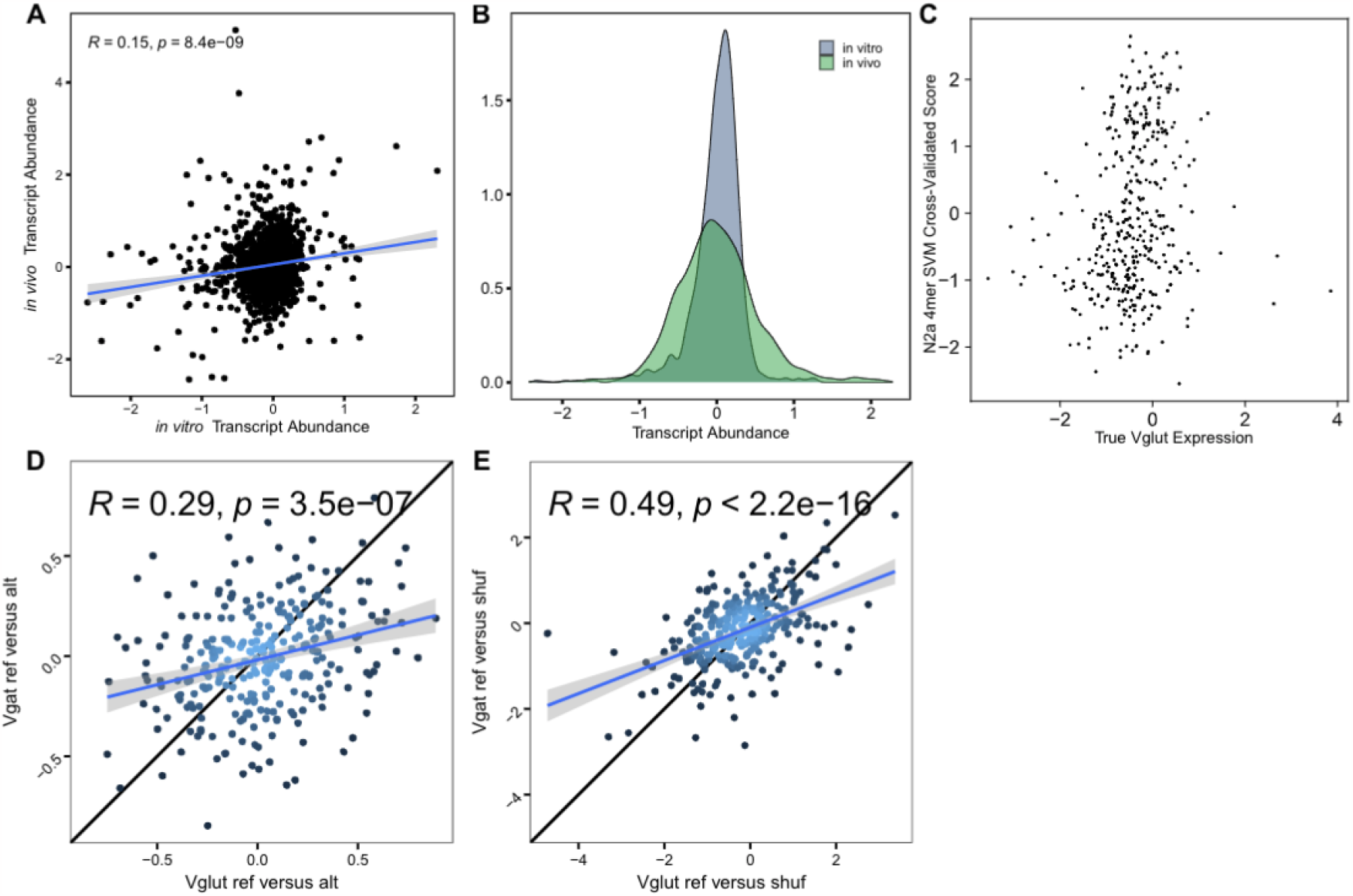
Low correlation and differences in range of expression between in vitro and in vivo suggest importance of regulatory context. A) Scatter plot of *in vitro* vs *in vivo* expression values show poor correlation. B) Comparison of *in vitro* and *in vivo* expression distributions, Brown-Forsythe Levene-type test for difference in variance p < 2.2e-16 C) N2a SVM score vs true Vglut expression show lack of correlation - indicating *in vitro* data cannot predict *in vivo* values. D) Scatter plot of *in vivo* VGAT and *in vivo* VGLUT expression values show better correlation.

### Comparison of effect sizes for barcodes relative to alleles

We next wanted to determine if our method was able to see smaller effect sizes that might be associated with single nucleotide variants, which comprise much of our variant set. One concern is barcodes - while the internal replication they provide can improve statistical power, it is possible that the effects of changing a 8-9 nucleotide barcode on transcript abundance may be more substantial than the 1 bp variant it tags. Including multiple barcodes per replicate is traditionally thought to allow correction for this by averaging or regression. Therefore, we modeled using random sets of barcodes drawn for the controls in a recent publication ^33^ to determine rates of false-positive ‘allelic effects’ driven by barcodes only [**Fig 5]**. We found that linear mixed modeling with 6 barcodes would be sufficient to remove the majority of barcode effects for most experiment designs, leaving a false positive rate of 8% **[Fig 5C]**, and generally small effects **[Fig 5A]**. This could be tolerable for many designs, however, if the design of the experiment is a screen for extremely rare functional effects (i.e., such as the rare ASD variants examined here), the false positive rate produced by 6 barcodes may be too high. Indeed, this is similar to the number of hits seen when testing for allelic effects here [e.g., 2-6 per assay, **Supp. Table 4]**, and thus we do not confidently report on the variant effects from this experiment. And while 1000’s of barcodes per variant can be added by PCR ^18,34,35^, the corresponding increase in library complexity would lead to substantial jackpotting in the cell-type specific *in vivo* context, where delivering each barcode to enough cells for robust measurement is a challenge. Thus, 6 barcodes might be sufficient for examining the large effects seen when measuring large effect size changes (e.g., the differences between elements) but more barcodes would be needed to assess smaller allelic effects, or when screening rare variants where only a small fraction are expected to be functional.

**Figure 5:**
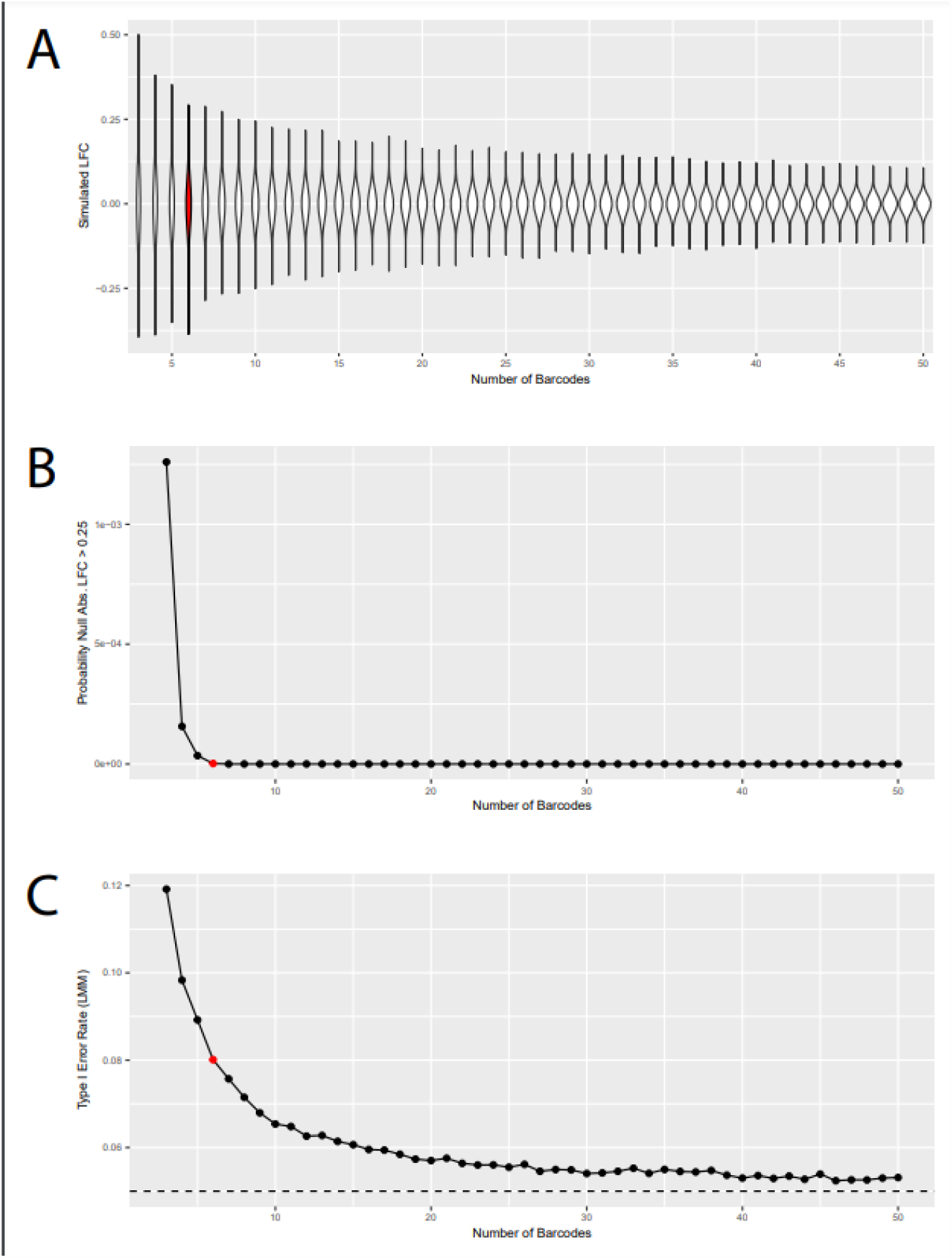
Simulated tests sampling control data to model barcode effects. Simulations drawing sets of barcodes for ‘allelic’ comparison using a barcode count of three through fifty for each allele (1,000,000 iterations, drawing from ∼100 Barcodes tagging the same control element from Mulvey et al, 2021). Red data points indicate six barcodes. A) Violin plots showing the range and frequency of log(2) fold-changes for randomly sampled data for the range of barcodes. B) Plot detailing the probability of obtaining a log(2) fold change less than or greater than 0.25 for randomly samples data for the range of barcodes. C) Plot detailing the Type I Error Rate of a linear mixed model for randomly sampled data for the range of barcode numbers.

## Discussion

Here we describe the development of a cell-type specific *in vivo* MPRA. We demonstrate that the method is sensitive enough to identify reproducible effects for hundreds of elements in parallel. It should be usable for dissecting the sequence dependence of previously identified regulatory elements with activity in neurons, using MPRA libraries designed for saturation mutagenesis of potential binding motifs ^18^ and other perturbations ^36^. It, with additional barcoding or other solutions, can also be adapted to examine allelic effects as well. Thus, the approach should have both translational and basic science applications.

This work and its challenges allowed us to deeply characterize the range of conditions, from environment to sequence context, that influence these regulatory assays. Our first attempts in delivering this library to rarer cell types using Rbp4 and Vgat Cre-lines were limited by low element recovery rates, making reproducible measurement of many variants in parallel intractable. Careful analysis of all stages of RNAseq library prep revealed jackpotting originated at cDNA synthesis, suggesting reporter mRNA was diluted beyond the point of efficient recovery. This is consistent with the relative sparseness of GABAergic cortical neurons compared to cortical excitatory neurons, indicating they will contribute less to the total RNA of the cortex. Use of neither emulsion PCR, reaction splitting, nor UMI incorporation in second strand synthesis could resolve this fundamental limitation. However, when we delivered to a more abundant cell type, increasing the barcode concentration in the final total RNA, the jackpotting was largely resolved. We do note that variability *in vivo* with AAV was still higher than when delivering to N2as in culture (PCC of >0.8 vs >0.9) with transfection, where delivering to >70% of cells at high copy number is straightforward. However, we were able to overcome this increased variability by increasing the sample number for *in vivo* assays. What other approaches might work to allow access to these rarer cell types and overcome the low barcode abundance in the starting RNA? Three general approaches come to mind: targeting AAV delivery to hit a larger portion of the Cre-positive cells (for example, here, we targeted the striatum, where a greater concentration of GABAergic cells exist), reducing the complexity of the library (using a smaller number of total barcodes, making each barcode more likely to be well represented), or enriching for the barcoded RNAs prior to cDNA synthesis, either by a targeted capture of reporter RNAs, or potentially purifying the Cre-positive cells by FACS or TRAP. Indeed, we did recently demonstrate the ability to measure allelic effects using TRAP in an enhancer context, especially when the library was delivered in a concentrated manner to a single anatomical structure so that library density is higher^37^. Any of these might further expand the current method to rarer Cre populations. Nonetheless, the current iteration of the technology already enables access to assessing elements in the regulatory context of mature cortical and striatal neurons, essential cell types for many CNS diseases. One key factor in the design, however, is the number of barcodes, as there is a clear tradeoff between adding more barcodes and elements which gains in experimental efficiency per animal, yet decreases reproducibility due to jackpotting, or barcode effects. For screening libraries where only a small fraction are expected to be functional, it may be preferred to reduce library complexity and barcode effects by removing barcodes entirely (letting the UTR serve as the barcode), or for designs that require barcodes (e.g., assessing enhancer variants) aim for ∼20 barcodes per allele **[Fig 5]**. Of course, libraries with larger effect sizes than SNPs (e.g, varying entire elements) do not need such a high level of stringency.

With regards to the role of these specific alleles in ASD, we are hesitant to make any specific claims since the number of variants showing significant effect was similar to what would be expected due to barcode effects in our simulation studies. Thus, we can’t be certain which allelic effects to confidently report. Nonetheless, there are two clear conclusions we can make. First, we can rule out large numbers of high-effect size UTR variants as a common cause for autism - if they were a common cause, we would have seen an excess of functional variants coming from the proband’s alleles relative to the unaffected sibling alleles, and we did not. Note - this does not rule out that in some rare instances a UTR mutation could be causal, but it will not be a frequent occurrence. Second, we can report that the typical effect sizes of single base mutations is small - even the significant ‘allelic’ effects detected were typically less than 25% changes, while differences in expression due to elements ranged over 8 fold. Thus, alleles with large functional effects will be rare.

Regardless, for elements of large effects, cell type-specific MPRAs *in vivo* for identification of functional elements across any Cre-defined cell type. This is important because the function of non-coding elements is strongly impacted by the suite of transcriptional regulators expressed in each cell type. Thus, this method presents the opportunity to perform these regulatory assays in the most relevant and specific biological context for a given disease. Altogether, we anticipate these methods will aid in the study of noncoding disease risk.

## Materials and Methods

### Animal Models

All procedures involving animals were approved by the Institutional Animal Care and Use Committee (IACUC) at Washington University in St. Louis, MO. Veterinary care and housing was provided by the veterinarians and veterinary technicians of Washington University School of Medicine under Dougherty lab’s approved IACUC protocol. All protocols involving animals were completed with: Tg(RBP4-cre)KL100Gsat/Mmcd (RRID:MMRRC_037128-UCD; Beltramo et al., 2013), Slc32a1tm2(cre)Lowl/J (catalog #16962, The Jackson Laboratory; RRID:IMSR_JAX:016962; Vong et al., 2011), and Vglut1-IRES2-Cre-D strain (Jackson Stock No: 023527). All mice were genotyped following a standard protocol of taking clipped toes into lysis buffer (0.5M Tris-HCl pH 8.8, 0.25M EDTA, 0.5% Tween-20, 4uL/mL of 600 U/mL Proteinase K enzyme) for 1 hour to overnight. This is followed by heat denaturation at 99 C for 10 minutes. 1 uL of the resulting lysate was used as a template for PCR with with 500 nM froward and reverse primers, using 1x Quickload Taq Mastermix (NEB) with the following cycling conditions: 94 1 min, (94 30 sec, 60 30 sec, 68 30 sec) x 30 cycles, 68 5 min, 10 hold.

### MPRA plasmid library preparation

For non-Cre-dependent reporter expression, we used a previously described pmrPTRE-AAV backbone which contained the following elements: CMV promoter, T7 promoter, mtdTomato CDS, hGH terminator, and flanking ITRs. The T7 promoter and mtdTomato CDS were amplified from pmrPTRE-AAV using PTRE_floxed_F/R and Phusion High-Fidelity PCR Master Mix (NEB). NotI and SalI sites added by the primers were used to subclone this amplicon into pRM1506_TMM432. The final pmrPTRE-floxed-AAV backbone consists of a floxed cassette containing the T7 promoter and tdTomato CDS in reverse orientation with respect to a CAG promoter, followed by a bGH terminator, all flanked by ITRs.

In order to determine if the elements in our library came from genes expressed in Vglut and Vgat regions specifically in excitatory neurons and medium spiny neurons we examined two single cell datasets ^38 39^GSE171977, and GSM4471659. The mean expression for each gene across all cells was calculated in both datasets, and ‘expressed’ was defined as exceeding the median of this value The data was then subsetted to only excitatory neurons or medium spiny neurons and the mean expression for each gene was calculated and expression was determined.. Genes were then converted back into their mouse orthologs using ensembl and these true false values were paired with their respective elements on table 1.

The specific oligo sequences with barcodes designed for this library are provided in **[Supp. Table 5]**. The UTR contexts for each oligo were taken from GRh37/hg19 by centering a maximum 120 bp window around the variant position. Variant allele sequences were substituted at the reference position to generate the alternative allele UTR context. For indels, the UTR context was limited to the minimum context that would fit either allele, and padding sequences were added outside of cloning cut sites. Additional elements with known or suspected post-transcriptional regulatory roles were included as well: the alpha component of the WHP posttranscriptional regulatory element (WPRE) and synthetic elements consisting of four tandem sequences for either the Smaug response element (SRE), Pumilio response element (PRE), or Quaking response element (QRE).

A constant 20 bp linker sequence separates the UTR context from a nine bp barcode sequence. Each UTR context was repeated in the design with six unique barcodes. Barcodes were selected to be Hamming distance of two apart and to exclude cut sites and homopolymers longer than three bases. Priming sites and cut sites were added to both ends to generate 210 bp oligos which were synthesized by Agilent Technologies.

The synthesized oligos were amplified with 4 cycles of PCR using Phusion polymerase and primers Bactin_FWD/REV. Amplicons were PAGE purified and digested with NheI and KpnI (NEB). Library inserts were cloned into pmrPTRE-floxed-AAV with T4 ligase (Enzymatics) and transformed into chemically competent DH5α (NEB). Outgrowths were plated on LB agar plates with 100 µg/mL carbenicillin, and approximately 71,000 colonies were counted, allowing us to capture the entire design at 95% confidence, assuming a 50% synthesis error rate. Plates were scraped, and the collected pellets were cultured for an additional 12 hours in LB with carbenicillin before preparing glycerol stocks and maxi preps (Qiagen).

### Cell culture

Mouse neuroblastoma N2a cells were maintained at 5% CO2 37°C, and 95% relative humidity in DMEM (Gibco) supplemented with 10% fetal bovine serum (FBS, Atlanta Biologicals). Human neuroblastoma SH-SY5Y cells were maintained similarly, except with DMEM/F12 (Gibco) substituted as the basal medium. Cells were also incubated with 1% penicillin-streptomycin (Gibco). For transient transfections, antibiotics were excluded from the transfection medium and re-introduced upon media change 12 hours post-transfection. Cells were passaged with 0.25% Trypsin-EDTA (Gibco) every 2-3 days or once they reached 80-90% confluency.

### Cell culture TRAP

For each cell culture TRAP experiment, six replicate T75 flasks (TPP or Sarstedt) were seeded in advance with mouse N2a neuroblastoma cells to be 80-90% confident by the time of transfection. For the library cloned into the pmrPTRE-AAV backbone, 20 µg of total plasmid containing an equimolar ratio of reporter library and an Ef1a-EGFP-RPl10a construct was transfected. For the Cre-inducible library, 23 µg of total plasmid was transfected, consisting of equimolar ratios of the library, an EF1a-DIO-EGFP-RPl10a construct, and an Ef1a-Cre construct. Transient transfections were performed with Lipofectamine 2000 (Invitrogen), and DNA:lipid complexes were prepared by co-incubation in Opti-MEM I (Gibco) for 30 minutes prior to transfection. Transfection medium was replaced 12 hours following transfection, and cells were harvested for TRAP after an additional 24 to 36 hours.

TRAP was performed as described ^24^ with minimal modification. Briefly, cells were incubated in 100µg/mL cycloheximide (Sigma) for 15 minutes at 37°C prior to harvest. Cells were rinsed twice with 5 mL of DMEM 100 µg/mL cycloheximide before being lifted into 5 mL of DMEM 100 µg/mL cycloheximide. Cells were pelleted by spinning at 500xg for 5 minutes at 4°C. The DMEM was replaced with 2 mL of ice-cold cell lysis buffer (10 mM pH 7.4 HEPES, 1% NP-40, 150 mM KCl, 10 mM MgCl2, 0.5 mM dithiothreitol, 100 μg/ml CHX, protease inhibitors, and RNase inhibitors) and cells were lysed on ice. Lysates were clarified by centrifugation at 2000xg for 10 minutes at 4°C. DHPC (Avanti) was added to a final concentration of 30mM, and lysates were incubated on ice for 5 minutes. A tenth of the volume was taken as the Input, and the remaining volume was incubated with protein L-conjugated magnetic beads (Invitrogen) coupled with a mixture of two monoclonal anti-GFP antibodies (Doyle et al., 2008). The beads were incubated for 4h at 4°C prior to four washes with a high-salt buffer (10 mM pH 7.4 HEPES, 1% NP-40, 350 mM KCl, 10 mM MgCl2, 0.5 mM dithiothreitol, 100 μg/ml CHX, protease inhibitors, and RNase inhibitors) before resuspension in cell-lysis buffer.

Input and TRAP RNA was extracted using Trizol LS (Life Technologies). Extracted RNA samples were DNase treated (Ambion) and cleaned by column-based purification (Zymo Research). Concentrations and RNA quality were determined using RNA ScreenTapes and a 4200 Tapestation System (Agilent Technologies). All RINe measurements exceeded 9.

Parallel Plasmid DNA for each replicate was recovered from each cell pellet following lysis using the Qiagen DNeasy Blood & Tissue Kit, and prepared for sequencing in parallel to RNA, as below. We found that having multiple replicate DNA libraries was critical for reducing variance in element activity measurements, at the transcript abundance level in particular. As such we recommend preparation of replicate DNA libraries, either from the plasmid input or from recovered plasmid from each experimental replicate of transfected cells.

### AAV9 vector production

The packaging cell line, HEK293, was maintained in Dulbecco’s modified Eagles medium (DMEM), supplemented with 5% fetal bovine serum (FBS), 100 units/ml penicillin, 100 mg/ml streptomycin in 37°C incubator with 5% CO2. The cells were plated at 30-40% confluence in CellSTACS (Corning, Tewksbury, MA) 24 h before transfection (70-80% confluence when transfection). 960 ug total DNA (286 ug of pAAV2/9, 448 ug of pHelper, 226 ug of AAV transfer plasmid) were transfected into HEK293 cells using polyethylenimine (PEI)-based method (Challis et al., 2019). The cells were incubated at 37°C for 3 days before harvesting. The cells were then lysed by three freeze/thaw cycles. The cell lysate was treated with 25 U/ml of Benzonaze at 37°C for 30 min and then purified by iodixanol gradient centrifugation. The eluate was washed 3 times with PBS containing 5% Sorbitol and concentrated with Vivaspin 20 100K concentrator (Sartorius Stedim, Bohemia, NY). Vector titer was determined by qPCR with primers and labeled probe targeting the ITR sequence (Aurnhammer et al., 2012). Titers used here ranged from 1 to 5 × 10^13^ vg/ml.

### *in vivo* MPRA

Two Vglut1-IRES2-Cre-D litters were subjected to intracranial injections for delivery of the library packaged in AAV9. P0-P2 pups were incubated on ice to anesthetize by inducing hypothermia for ∼10 minutes. An aliquot of the MPRA library packaged in AAV9 (∼10^9^ vg/uL) was drawn up in 33G Hamilton syringe with a 1 mm needle. Pups were brought up to the needle and 1 uL of virus was injected at three positions per brain hemisphere hemisphere (6 total injections per pup). Pups were taken directly to the warming pad until pups fully recovered (∼20 minutes). After recovery, pups were placed back into the cage with the mother and monitored every 24 hours for one week. At P21 brains were harvested for extraction of RNA.

We aimed to determine the source of this jackpotting and reasoned either the barcodes were all present in the starting template of total RNA and our library preparation was not efficient at a particular step, or the barcodes were simply too low abundance in the starting RNA pool. To this end, we conducted a series of technical replicates splitting a sample at each step of the library preparation protocol: cDNA synthesis, cDNA amplification, adapter ligation, and indexing PCR **[Supp. Fig 4A]**, and compared this to the low reproducibility observed between biological replicates [**Supp. Fig 4B]**. Taking a single RNA sample and doing two separate cDNA synthesis reactions for independent sequencing libraries resulted in jackpotting (PCC < 0.4) **[Supp. Fig 4C]**. Taking cDNA from a single sample and amplifying it in two independent reactions for library preparation also led to jackpotted samples (PCC < 0.4) **[Supp. Fig 4D]**. However, if the amplified cDNA from a single sample was taken into two independent reactions for adapter ligation, then the final sequencing libraries were highly correlated (PCC > 0.9) **[Supp. Fig 4E]**. This was the case for reactions split at the final indexing PCR as well (PCC > 0.9) **[Supp. Fig 4F]**. This result revealed to us that the source of jackpotting is at the cDNA synthesis or amplification steps. To investigate this further, we employed a variety of techniques that included reaction splitting/repooling to boost scale, unique molecular identifiers (UMIs), and emulsion PCR (ePCR). Reaction splitting/repooling and ePCR did not alleviate any of the jackpotting issues at any of the stages tested (data not shown). We then incorporated UMIs at the cDNA synthesis step in order to precisely quantify and eliminate PCR duplicates. After sequencing the resulting libraries and computationally collapsing UMIs, we found that only a small fraction of elements was recoverable. Together, these results led us to conclude that this jackpotting was, in fact, a representation of the barcodes present in the RNA: for a given amount pipetted (100 ng) from our total RNA from the brain, relatively few barcode molecules were present. Consistent with this, increasing input RNA up to 1 ug reduced jackpotting effects, but still resulted in relatively low sample correlations (PCC < 0.4). The possible solutions are to 1) increase the amount of RNA going into cDNA synthesis, which improved things to point (from PCC <.2 to PCC <.4) or 2) increase the library density by getting the library into a higher fraction of cells contributing to the total RNA.

### MPRA sequencing library preparation

Libraries were prepared by taking total RNA or TRAP RNA and performing cDNA synthesis using Superscript III Reverse Transcriptase standard protocol with pmrPTRE_floxed_AAV_antisense (GCATAAAAAACAGACTACATAATACTG) for library specific priming. Resulting cDNA or plasmid DNA, were then used for PCR to amplify libraries using Phusion polymerase (Thermo) using library specific primers pmrPTRE_AAV_sense (GCATGGACGAGCTGTACAAG) and pmrPTRE_floxed_AAV_antisense. Reactions were purified using AMPure XP beads between each step. The purified PCR products were then digested with NheI and KpnI restriction enzymes for 1 hour at 37 deg C. The purified digested products were ligated to 4 equimolar staggered adapters (this is to provide sequence diversity for sequencing). Ligated products were purified and then used for a second PCR using Illumina primers for library indexing. The purified libraries were then QC’ed and subjected to quality control and then 2×150 next generation sequencing on an Illumina NovaSeq.

### BC counting and normalization

Sequencing reads were trimmed using cutadapt v1.16 ^40^ and aligned to the library reference sequences using bowtie2 v2.3.5 ^41^ using “very sensitive” settings. Barcodes were counted from aligned reads with mapping quality of 10 or greater using a custom Python script. Counts within each sample were normalized to each sequencing library size using edgeR ^42^ as counts per million (CPM) prior to downstream analysis.

Measures of transcript abundance were calculated as the log_2_-transformation of the ratio of each barcode’s abundance, in CPM, from Input RNA over DNA, within each replicate. Barcodes with fewer than ten counts in either the RNA or DNA library were excluded from analysis. Similarly, ribosomal occupancy and translation efficiency were calculated by normalizing TRAP RNA to DNA counts and TRAP RNA to Input RNA counts, respectively. To calculate an element-wise measurement of transcript abundance or translation activity, a linear mixed effect model was fit which accounted for outlier barcode effects. Barcode-level measurements from each replicate were fitted to the formula Activity ∼ (1 | BC) using the lmer package in R where Activity may be either transcript abundance, ribosomal occupancy, or translation efficiency. The model intercepts were taken for each element as the summarised measure of activity.

### Element filtering and differential activity analysis

Differential effects of allele on transcript abundance and translation efficiency were each tested using a linear mixed effects model fitting random intercepts for barcode. This model was implemented using the lmer package in R with the formula Activity ∼ Allele + (1 | BC), where Activity may be either transcript abundance or translation efficiency. P-values were computed using a likelihood ratio test (LRT) with Activity ∼ (1 | BC) as the reduced model, and corrected for multiple comparisons by using the p.adjust function in R to apply the Benjamini-Hochberg procedure for false discovery rate.

Before testing, thresholds for element inclusion were determined by a grid search of count, barcode number, and replicate number thresholds that maximized the number of variants significant at a Benjamini-Hochberg FDR < 0.05. Briefly, at increasing count thresholds, variants within each replicate were retained if both alleles had more than a set threshold for barcodes above said count threshold, and variants with both alleles passing count and subsequently barcode thresholds in a minimum number of replicates were selected for analysis. An arbitrary count threshold minimum of 10 counts was enforced. For the *in vitro* MPRA, variants must be present with both alleles having three barcodes with at least 10 counts from both RNA and DNA in at least four replicates. For the *in vivo* MPRA, variants must be present with both alleles having five barcodes with at least 10 counts from both RNA and DNA in six replicates.

### Fluorescent immunohistochemistry and analysis

Brains were harvested from postnatal day 21 mice, and one hemisphere was chosen for subsequent RNA extraction/TRAP. The remaining hemisphere was fixed for 48 h in 4% paraformaldehyde followed by 24 h in 15% sucrose in 1× PBS and then 24 h in 30% sucrose in 1× PBS. The hemisphere was then frozen in OCT compound (optimum cutting temperature compound; catalog #23-730-571, Thermo Fisher Scientific). A Leica CM1950 cryostat was used to create 40 μm sagittal sections of brain tissue. Sections were immediately placed in a 12-well plate containing 1X PBS and 0.1% w/v sodium azide.

For immunostaining, sections were incubated in a blocking solution (1× PBS, 5% donkey serum, 0.25% Triton-X 100) for 1 h in a 12 well plate at room temperature, then with rabbit anti-RFP primary antibody (1:500; Rockland catalog #600-401-379) in blocking solution overnight in a sealed 12 well plate at 4°C. Following three five-minute washes in PBS, sections were incubated in donkey anti-rabbit Alexa Fluor 568 secondary antibody (1:1000, Invitrogen catalog #A10042) and DAPI (in blocking solution for 1 h. Sections were washed as before, and during the second wash, 1 μg/mL DAPI was added. Sections were slide mounted with Prolong Gold and visualized for anti-RFP and DAPI staining on a Zeiss Axio Imager Z2 four-color inverted confocal microscope. TdTomato-positive cells were quantified by hand using FIJI ^43^.

### Machine learning

Gapped k-mer SVM models were fit using gkmSVM ^25^ with the parameters -l 4 -k 4 -m 1 (4-mers) and -l 5 -k 5 -m 1 (5-mers). Stratified five-fold cross-validation and computing ROC and PR curves was performed using scikit-learn version 0.19.1 ^44^.

### Data and reagent availability

MPRA libraries are available upon request. MPRA data are deposited with GEO at GSE186455, to be made public upon publication. Reviewer token for peer review is: uxgduikajbijpkz. Code is available at bitbucket: https://bitbucket.org/jdlabteam/mpra_lib_1.0_methods_paper/src/master/

## Acknowledgements

We’d like to thank Bernie Mulvey, Dana King, and the Djuranovic lab for discussions and advice. We’d also like to thank Kristian Quigless, Christian Doss, as well as Mingje Li and the Hope Center Viral Vectors Core for technical support, as well as the CGS spike-in cooperative (especially Jess Hoisington-Lopez and MariaLynn Crosby), and GTAC@MGI for sequencing support. This work was funded by the Simons Foundation (571009) and the NIH (5R01MH116999) and T32 (MH014677, GM007067).

## Glossary

AAV: adeno-associated virus
ASD: Autism spectrum disorder
AUPRC: area under the precision recall curve
AUROC: area under the receiver operating characteristic
BC: Barcode
CNS: Central nervous sytem
DiO: Double-floxed inverse orientation
MPRA: Massively Parallel Reporter Assay
PRE: Pumilio response element
QRE: Quaking response element
RBP: RNA binding protein
SRE: Smaug response element
SVM: Support vector machines
TE: Translation Efficiency
TF: Transcription factor
TRAP: Translating Ribosome Affinity Purification
UTR: Untranslated Region
WPRE: WHP posttranscriptional regulatory element
FDR: False discovery rate
LFC: Log fold change
LRT: Likelihood ratio test
PCC: Pearson’s Correlation Coefficient

## Figures

**Supplemental Table 1: List of variants used for MPRA library, and corresponding transcripts**. *SeqName* - concatenation of gene name and sequencing data collection (either Whole Genome Sequences, or SSC exome sequencing) the sequence came from. *Hg38positionTag -* location and mutation, in hg38 genome build, along with family ID. *Pheno* - was mutation from case or sibling control. *FamilyID, SampleID* - for individual, from SSC. *ensembl_gene_id* - Ensembl gene identifier, *tx_id* - corresponding transcript identifier. Mutations hitting more than one transcript have multiple rows. *Library_seq_ref* - the oligonucleotide sequence for the reference allele, *Library_seq_alt* - the oligonucleotide sequence for the alternative (mutant) allele. *Library_seq_shuf* - the oligonucleotide sequence generated by random shuffling of the reference sequence (done for 50% of ref seqs). *Oe_lof_upper (aka LOEUF)*: a measure of constraint for this gene, derived from GNOMAD database ^45^. *Sc_ne_expression_higher_then_median* and *sc_ms_expression_higher_then_median*: True/False values for if transcripts of this gene were expressed (cellwise mean CPM of a single transcript greater than or equal to geneswise median mean CPM) for excitatory neurons from mouse cortex single cell data ^39^, specifically GEO accession GSM4471659, and for medium spiny neurons from mouse single cell striatum data specifically GSE171977, respectively.

**Supplemental Table 2: Overview of replicates, sequencing statistics, and results for all MPRA experiments**. Column headers and abbreviations: *pDNA* - plasmid DNA, *vDNA* - viral DNA, *Read depth per replicate (range)* - the amount of usable sequences (after alignment and QC, per replicate) *Cor across replicates*- The pairwise correlations between transcript abundance(RNA/DNA) for all elements across biological replicates. Range of correlations provided. *% of barcodes detected* - % of barcodes from the starting oligonucleotide library detected. *Counts per barcode* - Average counts per barcode. *Number hits (ref v shuf) [Abundance]* - number of elements showing significant (FDR <.1) and at least 25% magnitude (LogFC +/- .32).

**Supplemental Table 3**: **Log**_**2**_ **Fold-Change of Reference vs. Shuffled Elements**. Worksheet detailing the results of linear mixed modeling in different cell contexts. TE prefix on sheets denotes translational efficiency analysis from TRAP experiment, all others are mRNA expression differences. Results shown include log_2_ fold-change of the reference element vs. the shuffled element, the associated p-value, Benjamini-Hochberg style false discovery rate and Bonferroni multiple test corrections, the patient phenotype (ASD case or not), SFARI gene score, LOEUF value from the gnomAD database, constrained column (LOEUF < 0.268), and whether the gene was found to be expressed in excitatory neurons and medium spiny neurons. N2A 410 denotes Neuro-2a cells co-transfected with the barcoded library and Cre recombinase vector experiment, Vglut 410 denotes the Vglut1-IRES-Cre animals injected with viral barcoded library experiment, and Vgat 410 denotes the Vgat-IRES-Cre animals injected with viral barcoded library experiment. Please refer to the Animal Models section of Materials and Methods for Jackson catalog numbers and further information. Elements not passing QC cutoffs for statistical analysis based on read depth/representation are denoted with “Not_Tested” in the logFC columns.

**Supplemental Table 4**: **Log**_**2**_ **Fold-Change of Reference vs. Variant Elements**. The same layout as Supplemental Table 3, but now comparing the reference element vs. the variant (alt) element identified in the patient allele.

**Supplemental Table 5**: Oligonucleotide sequences included in the MPRA library and their corresponding barcodes.

## Supplemental

**Supplemental Figure 1.**
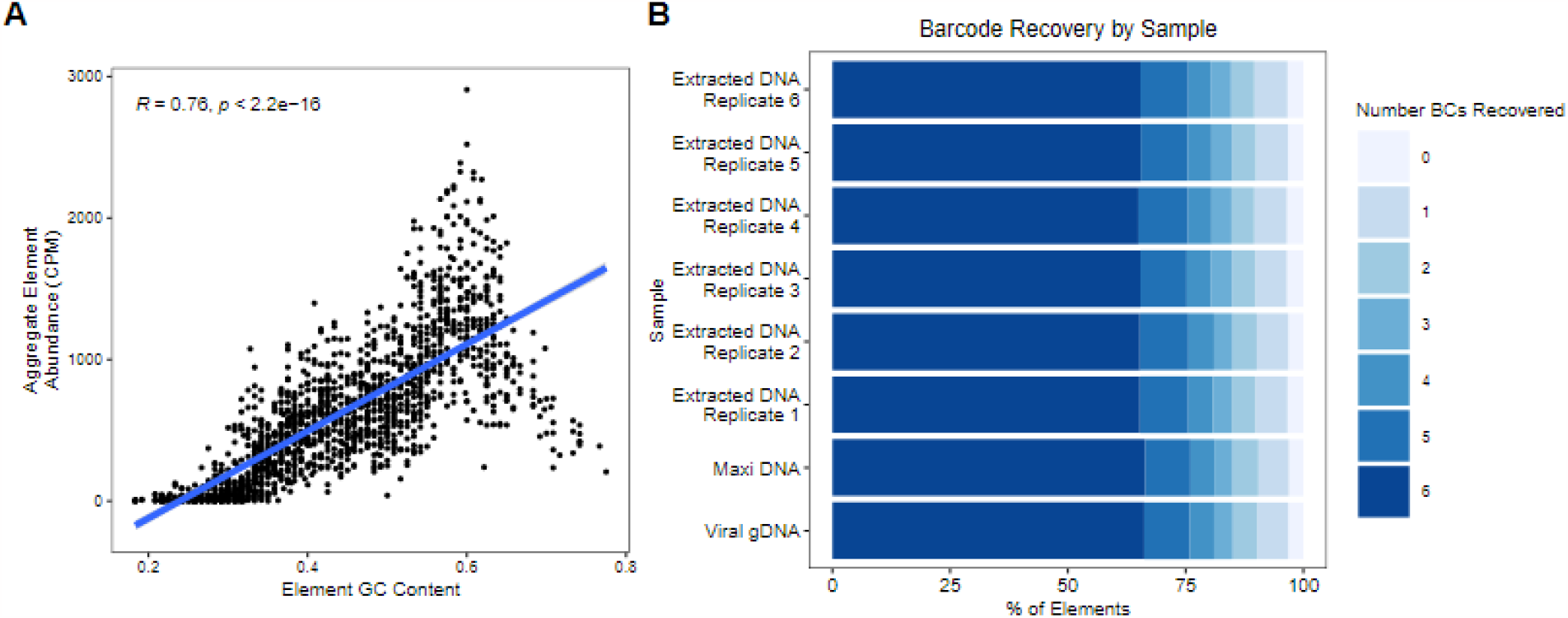
Library quality control. A) Average element abundance vs element GC content B) Barcode recovery by sample

**Supplemental Figure 2.**
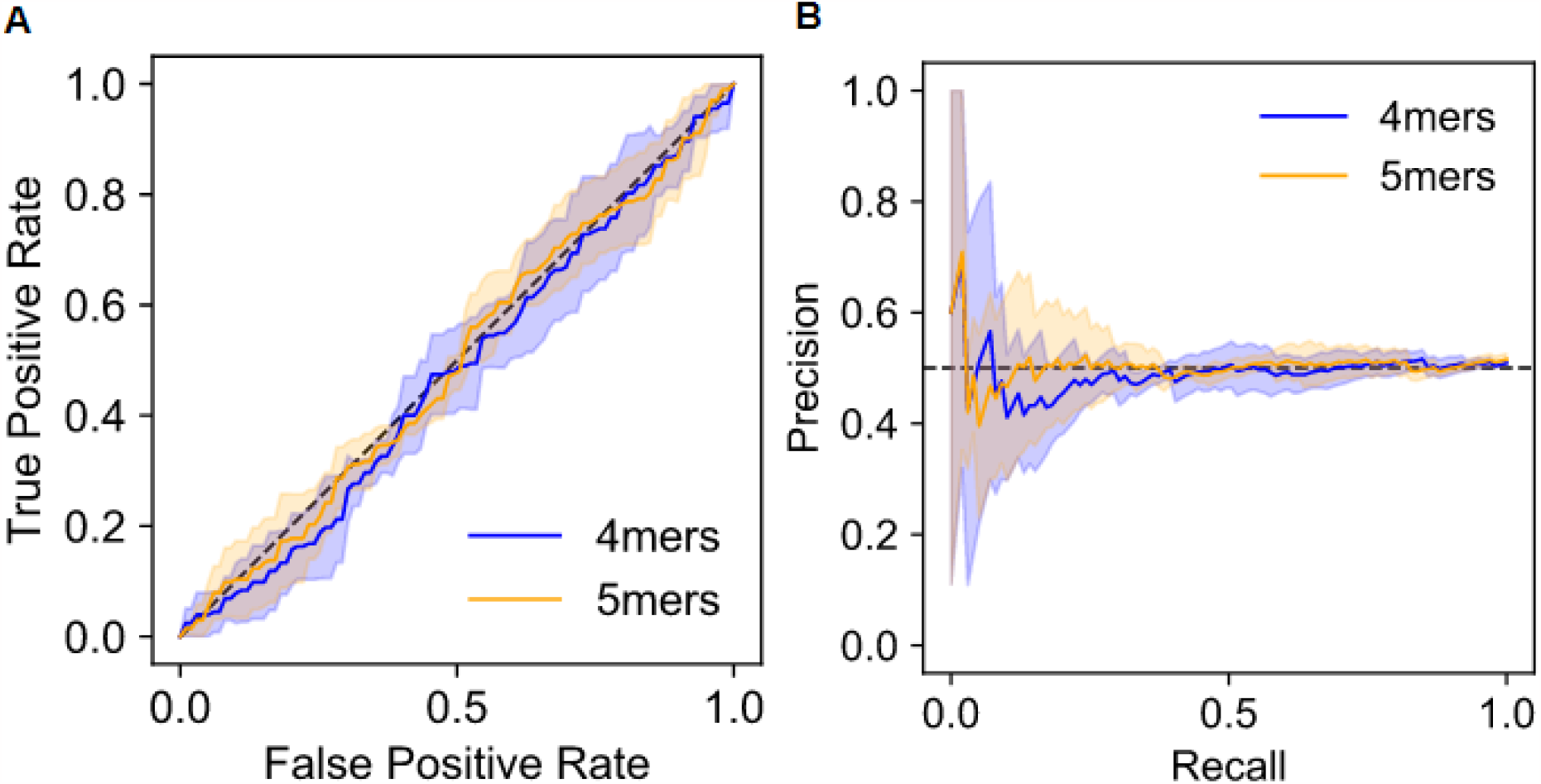
SVM data from random labels. A) ROC and B)PRC for k-mer SVMs to classify high and low expressing shuffled elements. Shaded area represents 1 standard deviation based on five-fold cross-validation

**Supplemental Figure 3.**
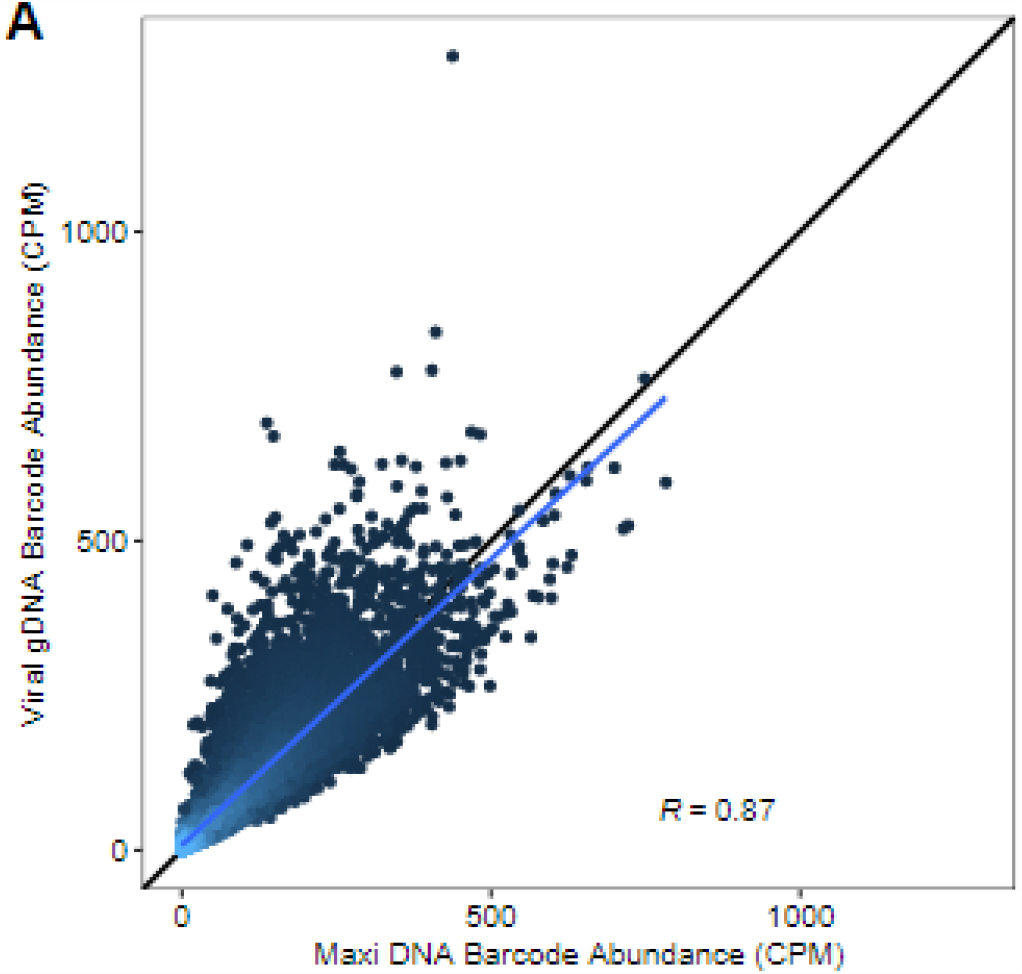
Viral packaging correlates with plasmid DNA barcode counts. A) Scatter plot showing correlation between plasmid DNA and viral DNA barcode counts

**Supplementary Figure 4:**
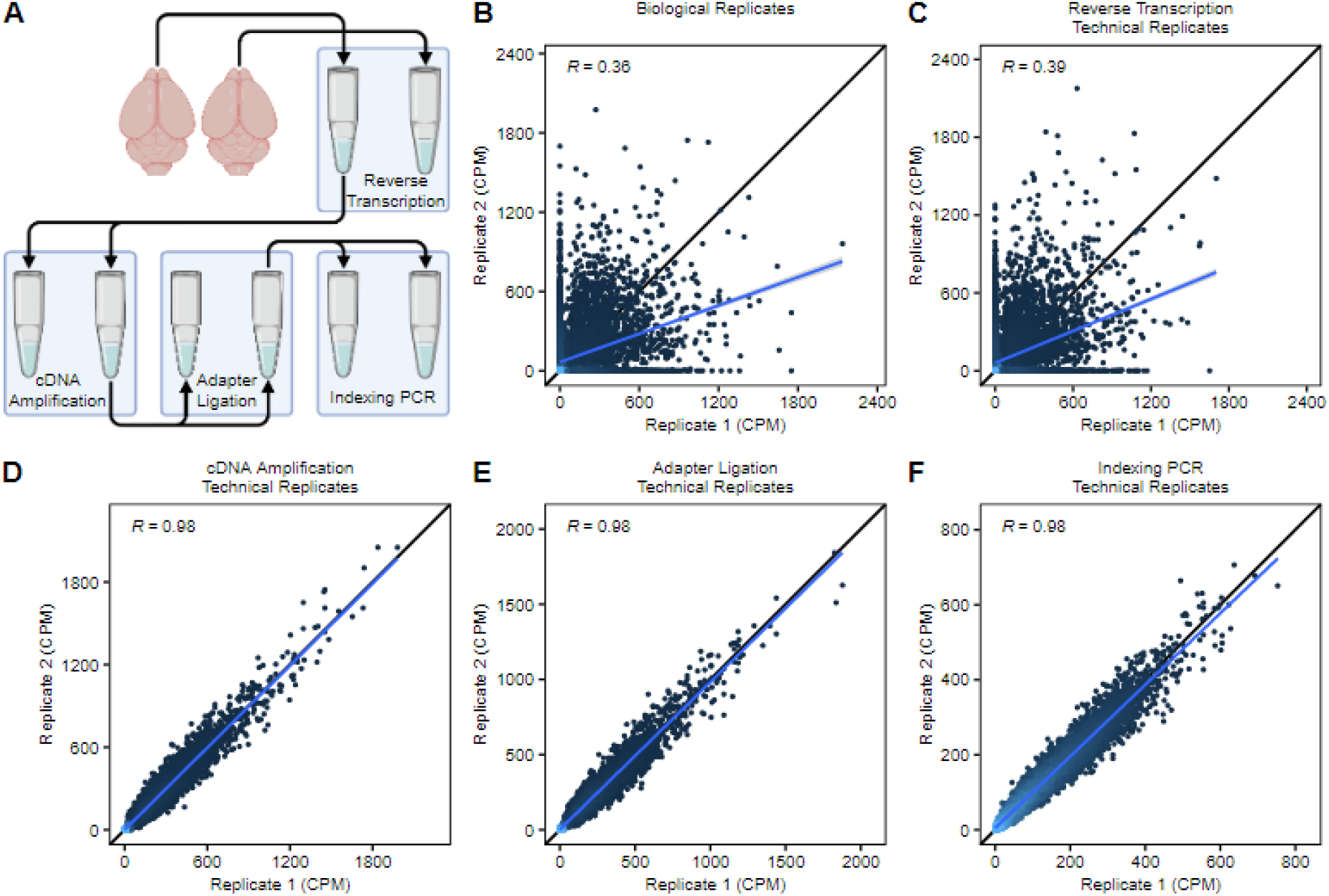
Reaction splitting to determine source of jackpotting. A) Various steps of the MPRA library preparation pipeline. Brains are harvested and extracted total RNA reverse transcribed to create cDNA, cDNA is amplified, adapters are ligated on, and sequencing indexes are added to complete libraries. B) Representative scatter plot showing correlation of biological replicates from Rbp4/Vglut animals C) Library correlation of technical replicates when splitting at the cDNA synthesis stage. D) Library correlation of technical replicates when splitting at the cDNA amplification stage. E) Library correlation of technical replicates when splitting at the adapter ligation stage. F) Library correlation of technical replicates when splitting at the indexing PCR stage.

